# Evaluation of the fluid-movement contribution to the oxygen transport in tissue

**DOI:** 10.1101/2021.01.28.427630

**Authors:** Zhu Liu, Chenyu Wen, Shi-Li Zhang

## Abstract

The model of diffusive O_2_ transport from capillary to tissue was established by Krogh a century ago. This model is incomplete as it ignores the often inevitable convective O_2_ transport via fluid movement. Here, we propose a one-dimensional physical-phenomenological model to evaluate the contribution of fluid movement to the O_2_ transport in tissue. Both the O_2_ gradient and the total O_2_ flux are found to be sensitive to the fluid movement. For small flow rates with a Peclet number *Pe* < 1, a critical flow rate, *u*_dc_, is introduced to characterize the contribution of fluid movement to the O_2_ transport, as well as to evaluate the fluid contribution in O_2_-deficient tissues and the cytoplasm movement inside muscle fibers. During hemostasis, the O_2_ flux contributed by the interstitial flow even below a rate of 2 μm/s, although negligible near the capillary, can be significant for the tissue residing far from the capillary. For an isolated intramyocyte mitochondrion, the cytoplasm movement can play a key role in the O_2_ transport. These considerations point to the scenario of an external force to drive the fluid movement leading to an accelerated O_2_ transport to sustain the mitochondrial consumption. Our model offers a comprehensive picture of the O_2_ transport in tissue by including both the concentration gradient and the hydraulic pressure gradient. It predicts that even with a small external force, damaged tissues with higher permeability can yield larger rates of interstitial flow promoting O_2_ transport for tissue recovery.

**Highlights:**

1. The fluid movement can play a significant role in O_2_ transport in tissue with high Prandtl number *ν/D* ~ 10^3^;
2. A critical flow rate, *u*_dc_, is introduced to evaluate the convective transport flux by the fluid in tissue for Pe < 1.
3. O_2_ transport in tissue is a result of the balance between hydraulic pressure gradient induced flux and concentration gradient induced flux.
4. Our model indicates more O_2_ pumped by convective transport into the O_2_ -deficient region where the O_2_ diffusive flux cannot penetrate.
5. The cytoplasm movement can excite more O_2_ flux to the mitochondria with higher myoglobin concentration.

## 1. Introduction

Human bodies rely on oxygen metabolism in mitochondria to produce the cellular energy adenosine triphosphate (ATP). During respiration, O_2_ is taken up by hemoglobin forming Fe-O_2_ bonds inside red blood cells. It is, then, circulated through blood streams to capillaries in peripherals. In capillaries, O_2_ dissociates from hemoglobin, dissolves in plasma, and passes through the endothelium of capillaries. Finally, it reaches tissue through the interstitial flow and transports to mitochondria so as to participate in the ATP production. From capillary to tissue, O_2_ confronts the highest transport resistance due to its low solubility in the body fluid [1, 2].

Krogh [3] introduced a model to describe the oxygen transport from capillary to tissue based on the passive diffusion of O_2_ inside the tissue. Since then, the subsequent models have been developed in different scales from tissue to organs with various geometries in multiple types of muscles and tumors [4]. Similarly, all these models are constrained to pure diffusive transport, driven by concentration gradient, without consideration of the O_2_ convective transport originating primarily from the fluid movement. Recently, experimental data support a paradigm shift away from merely counting on the Krogh’s diffusion theory [5]. 1) Several experiments have demonstrated that the O_2_ concentration gradient (Δ*P*_O2_) of intramyocytes is negligible during both the rest and exercise [6, 7], which contradicts the pure diffusion model that implies a varied O_2_ concentration with distance; 2) Δ*P*_O2_ between the microvascular and the interstitial space during the contraction was almost unchanged compared to that in rest [8, 9]; 3) Mass balance analysis for O_2_ transport shows that the O_2_ permeability (Krogh constant, *K* = *Dα*) *in vivo* is 1-2 orders of magnitude greater than *in vitro* value [10]. The further scrutiny of *in vivo* and *in vitro* experimental data suggests that the microvessel wall consumption has a negligible contribution to the O_2_ flux *in vivo* [11]. Hence, it is crucial to reassess the transport through fluid movement via interstitial space from microvessel to tissue.

The O_2_ transport from the capillary to the tissue is conveyed by the interstitial fluid, a dynamic movement flow for maintaining the nutrition metabolism of body, which plays a crucial role for the O_2_ transport from the capillary to the tissue mitochondria [12]. In the interstitial fluid, the Prandtl number, *ν/D*, of O_2_ is ~10^3^, where, *ν* = 10^−4^ cm^2^/s is the kinematic viscosity of interstitial fluid and *D*_O2_ = 1.6×10^−2^ cm^2^/s is the O_2_ diffusion coefficient. The large Prandel number indicates that convective transport in the body flow of the interstitial fluid is more effective than diffusive transport even at low flow rates [13].

The interstitial flow is a one-way movement from the capillary to the initial lymphatics. The interstitial fluid includes blood plasma, water, proteins, and electrolytes leaked from capillaries [14]. For healthy tissue, 10-20 % of the excess interstitial fluid is drained by the initial lymphatics and the rest is reabsorbed by venous capillaries. Hence, normally, the lymph formation and the lymph flow rate regulate the flow rate of interstitial fluid. Lymphatic vessels locate inside the connective tissues and distribute across the entire muscle tissues correlated with arcading arterioles and veins [15]. In muscle tissues, the initial lymphatics has a luminal size about 10-100 μm with an average distance of several hundred micrometers [15, 16]. However, the initial lymphatics does not have smooth muscles, and thus cannot contract spontaneously. Their openness is controlled by the contraction of surrounding muscle fibers and arterial pulsations [15, 17]. The typical interstitial flow rate, *u*, is in the range of 0.1 to 2 μm/s [18]. During exercises, the lymph flow can increase by more than 10-30 folds [14, 19]. During lymphedema, the lymph flow is almost ceased due to disease or injury, rendering the interstitial fluid and tissue swollen [16, 20]. Therefore, the interstitial flow maintains the nutrition and body fluid metabolism. The entire process is balanced by the venous pressure, interstitium composition, interstitial pressure and pressure inside lymphatic vessels [18, 21].

The interstitial flow through tissue has been found to facilitate protein transport, drug deliveries, and therapeutics [22, 23]. Convection is an active process prone to local external force offered by, such as, massage and body stretch [14, 24]. The interstitium can tune its porosity and permeability responding to tissue deformation and compression [25, 26]. Furthermore, manipulating large molecules through interstitium to tumor sites by enhancing its convective transport has been investigated as an effective means in cancer therapies [27, 28]. In addition, the convective transport of interstitial flow is important in nutrition delivery for cartilage, bone modeling[29, 30], and soft tissue [18]. The morphogenesis, differentiation, and regeneration of cells are closely associated with the interstitial flow [18, 31–33]. Moreover, the interstitial flow has been correlated to the meridian channel theory [34, 35]. The significance of the convection transport by the interstitial flow is tied to the molecular size [36]. Normally, diffusion dominates for small size molecules, such as O_2_, and short distance transport. But if the tissue is under the condition of impaired capillaries and cannot sustain with an effective Δ*P*_O2_, the convective transport through the flow inside the tissue cannot be ignored.

Another Peclet (*Pe*) number of the flow (*ub*/*D*) has also been used to evaluate the convection transport of the fluid movement contribution, where *b* is the tissue scale. Typically, the diffusion contribution dominates for *Pe* < 1 and the convection dominates for *Pe* > 1. Specifically, *b* is around 100 μm for typical tissue, such as the muscles, *Pe* is around 0.1 at a typical interstitial flow rate of 1 μm/s. As a result, the fluid movement contribution to the O_2_ transport is considered negligible. Δ*P*_O2_ can also affect the contribution evaluation of the convection [13]. Moreover, the continuity of interstitial spaces across tissue and organ boundaries in human has recently been confirmed [37]. Hence, the consumption flux should be included in the convective contribution, since the local diffusion gradient is correlated to the consumption rate in tissue flux.

In this work, we evaluate the fluid movement contribution to the O_2_ transport in tissue. The convective diffusion transport process is thoroughly solved in a one-dimensional (1D) framework by introducing a critical flow rate, *u*_dc_, and a spatial convection-diffusion boundary, *x*_0_. First, the convective-diffusion transport equation is solved by applying the boundary conditions of *C*_O2_ = *C*_0_ at *x* = 0 and ∇*C*_O2_ = 0 at *x* = *b*. In addition, the convection contribution is evaluated by the fluid movement inside the muscle tissue with different myoglobin concentrations. Darcy’s law is introduced to relate tissue permeability to local pressure gradient. A comprehensive picture of O_2_ transport in tissue is proposed by considering the convective component of the flow and the diffusive component driven by concentration gradient. Be noticed here, a different terminology of “advective transport” has been used in different literature to describe the convective transport of the fluid movement.

## 2. Methods and modeling

The fluid movement in tissue represents a dynamic flow including cytoplasmic streaming and interstitial flow. Here, a homogenous media is used to describe the O_2_ transport in the tissue. The O_2_ flux in the tissues at any site, *J*_total_, includes two parts: molecular diffusion in the stasis fluid, *J*_diff_, and convection of fluid movement, *J*_conv_, *i.e*.:

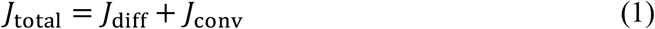

The O_2_ diffusion across a surface of unit area per unit time is described by Fick’s first law:

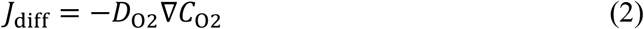

where *C*_O2_ is O_2_ concentration determined by its partial pressure (O_2_ tension) *P*_O2_ via *C*_O2_ = *αP*_O2_, with *α* being the O_2_ solubility coefficient in the body fluid. The convective transport by the fluid flow at a flow rate *u* is [38]:

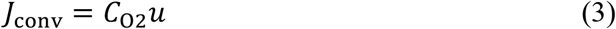

Assuming a homogeneous consumption of O_2_ by mitochondria inside the tissue at a rate of *M*_0_, the O_2_ transport in a volume tissue is equal to the O_2_ flux through the tissue boundary and the consumption within the tissue volume, which can be described by Fick’s second law:

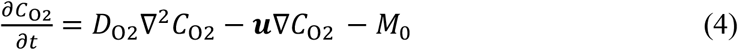

For an impressible fluid, ∇***u*** = 0. At steady state, *∂C*_O2_/*∂t* = 0, and the above equation becomes:

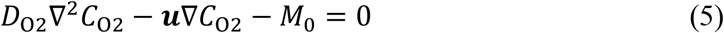

Without the convection term ***u***∇*C*_O2_, Eq. 5 is reduced to Krogh’s model that uses a cylindrical coordinate to solve the O_2_ diffusion radially from the capillary to the muscle tissue with the capillary centered.

The fluid movement contribution to the O_2_ transport is evaluated as follows. It is difficult to analytically solve Eq. 5 in two-dimensional (2D) or three-dimensional (3D) domains. It can be solved analytically in the 1D domain with the boundary conditions of *C*_O2_ = *C*_0_ = *αP*_0_ at *x* = 0 where the O_2_ source is located and ∇*C*_O2_ = 0 at *x* = *b* where the O_2_ diffusion cannot penetrate through. The close form of the solution to Eq. 5 is:

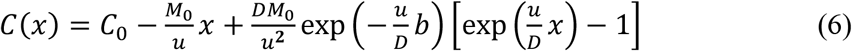

where, *D*_O2_ = *D*. Substituting Eq. 6 into Eq. 1-3, *J*_diff_ and *J*_conv_ can be obtained as follows:

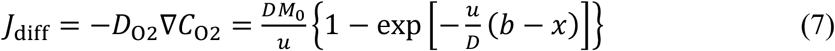

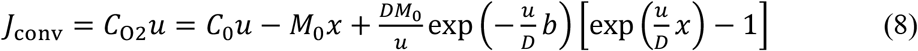

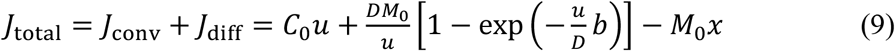

Details in mathematical derivations can be found in the supporting information. The distribution of O_2_ concentration along *x* can be resolved, so does the convective and diffusive transport fluxes.

In addition, a 1D numerical illustration is implemented using COMSOL Multiphysics in Fig. 1. The flux into the tissue at *x* = 0 and *u* = 0 is determined by the total O_2_ consumption flux, *J*_m_. Here, the illustration is performed with *M*_0_ = 2.3 ×10^−5^ ml O_2_/ml s, *b* = 1 mm with *J*_m_ = *bM*_0_ = 2.2×10^−7^ ml O_2_/cm^2^ s. The cytoplasm movement inside the muscle fiber is evaluated by changing the myoglobin concentration. The correlation of the flow rate *u* to the tissue permeability is studied by referring to Darcy’s law.

**Fig. 1.**
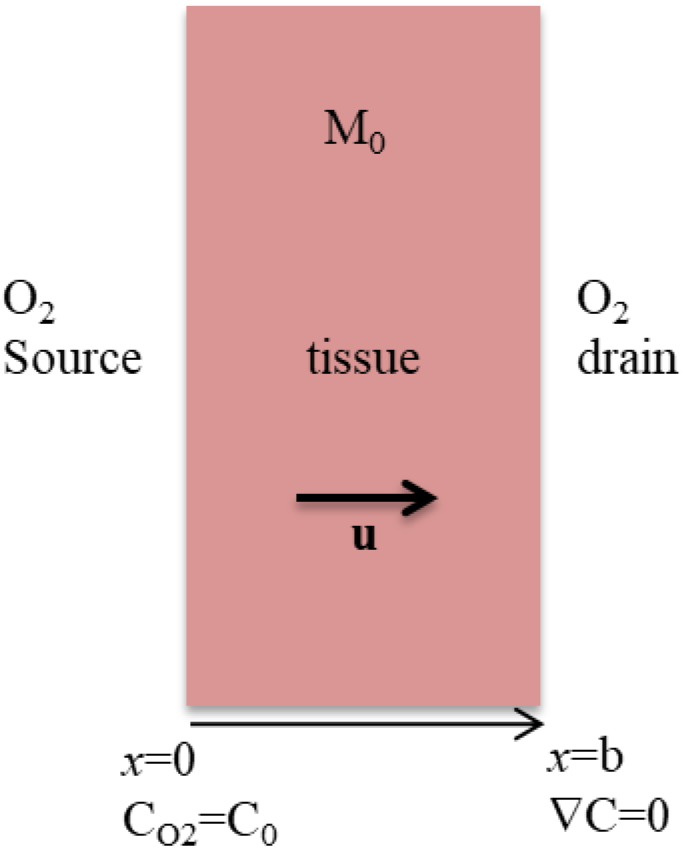
Illustration of a 1D model for the O_2_ transport.

## 3. Results and discussion

The numerical analysis starts from a simple situation without consideration of fluid movement, *i.e*., convention velocity *u* = 0. The *P*_O2_ distribution across the tissue is shown in Fig. 2(A). The penetration length of O_2_ is determined by *L*_p_ = (2*DC*_0_/*M*_0_)^1/2^ [4] (see supporting information) at *u* = 0, where *J*_diff_ = 0, *i.e*., ∇*C*_O2_ = 0 at *x* = *L*_p_. Thus, *L*_p_ measures the maximum distance at which O_2_ can be delivered solely by diffusion. –In this situation, part of the tissue is under the condition of hypoxia and fatigue. The critical initial *P*_O2_ with *b* = *L*_p_ is *P*_O2_(*L*_p_) = *b*^2^*M*_0_/(2*DαC*_0_).

**Fig. 2.**
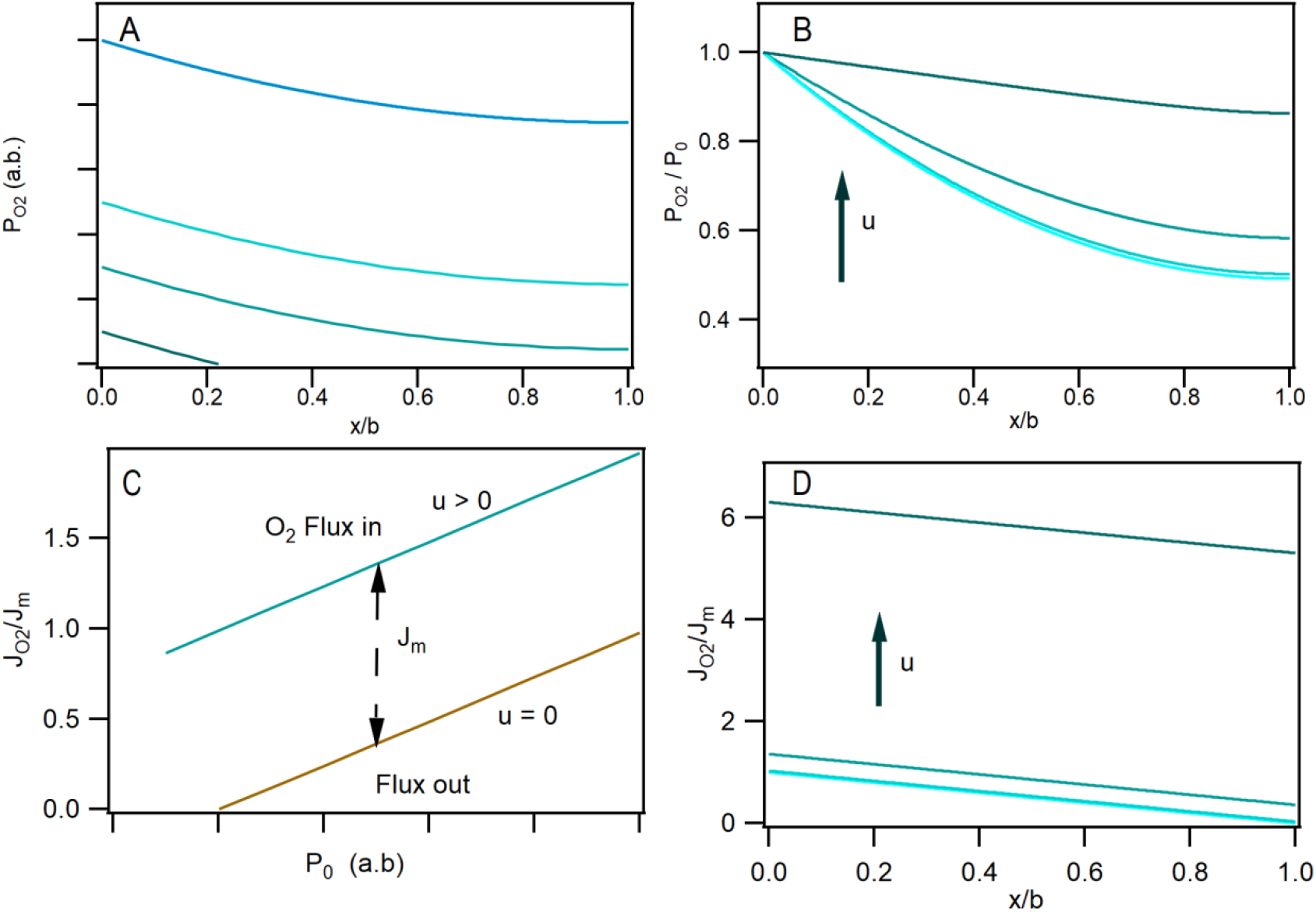
Concentration profile at different (A) O_2_ tension of capillary; (B) fluid flow rate and (D) normalized O_2_ flux distribution cross the tissue (0, 0.1, 1 and 0.1 μm/s) at PO_2_ = 50 mmHg (*J*_m_= 2× 10^−7^ ml O_2_ /cm^2^ s); (C) the O_2_ flux in (x=0) and the flux out (x=b) as the function of P_O2_.

By introducing the convective transport (*u* > 0), the picture is changed. As *u* increases, ∇*P*_O2_ decreases, while the O_2_ concentration at *x* = *b* increases, as shown in Fig. 2(B). The difference in O_2_ flux between in and out the tissue is determined by *J*_m_, as shown in Fig. 2(C). Moreover, the corresponding O_2_ flux, *J*_O2_, shows an enhanced O_2_ flux into the tissue (*x* = 0) as *u* increases, which is beyond the consumption by the tissue from Fig. 2(D). The *J*_O2_ across the tissue *b* is decreased due to the O_2_ consumption. As *u* increases, higher *J*_O2_ can penetrate through *x* = *b*, even though ∇*P*_O2_ cannot reach at *u* = 0. It suggests that the O_2_ flux through the tissue would not only be constrained by the local *J*_m_ but also the flow rate *u*.

The normalized O_2_ flux at *x* = 0 is used to characterize the total O_2_ injected into the entire tissue region *J*_total_ (from 0 to *b*) as shown in Fig. 3(A). At very low flow rate, *J*_total_ remains almost unchanged. However, it grows exponentially when the flow rate increases therefrom. *J*_conv_/*J*_total_ is displayed in Fig. 3(B). Three regions of O_2_ transport can be discerned: i) convection wherein the O_2_ transport is predominated by the fluid movement, and *J*_total_ is linearly dependent on *u*; ii) mixed convection-diffusion in which both *u* and Δ*P*_O2_ have large effects on the O_2_ transport and *J*_total_ increases dramatically with *u*; and iii) diffusion where *u* is too small to affect Δ*P*_O2_ and the O_2_ transport is dominated by diffusion. In general, the convection (*J*_conv_) and diffusion (*J*_diff_) contributions can be evaluated by *Pe*.

**Fig. 3.**
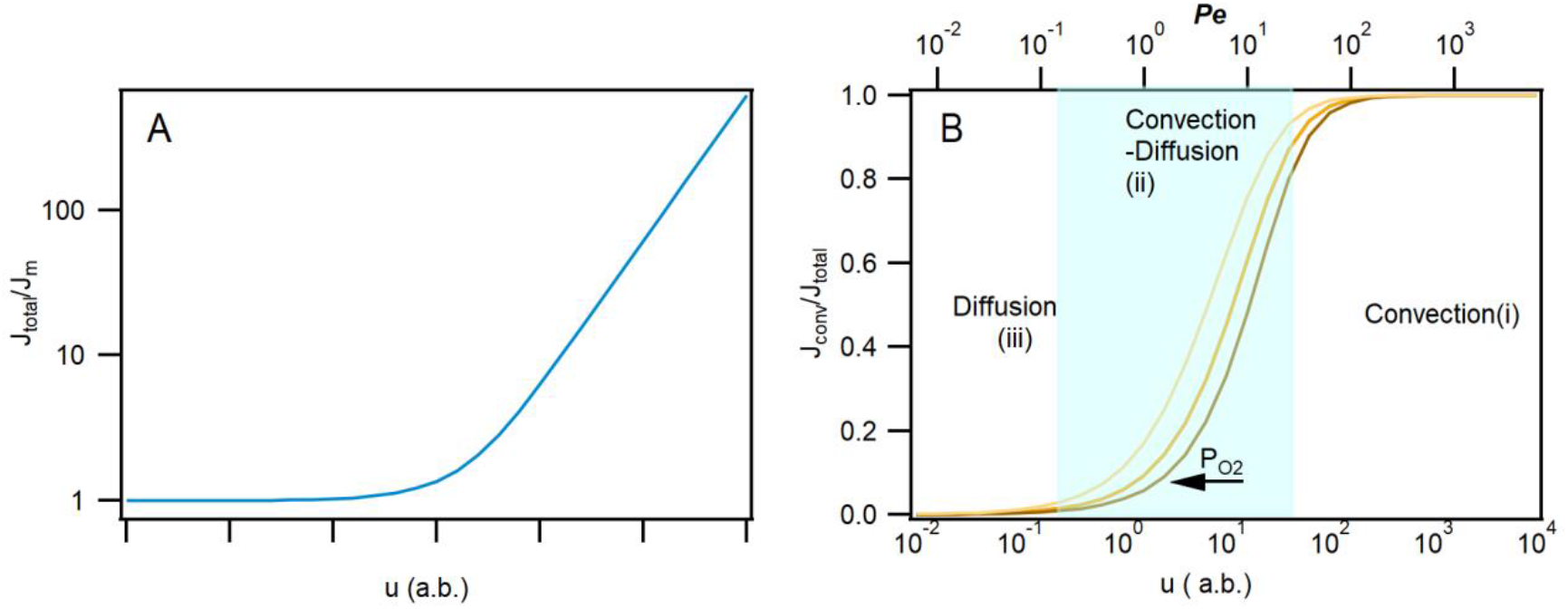
At x = 0, (A) J_total_/*J*_con_ at *P*_O2_ = 50 mmHg (*J*_m_= 2× 10^−7^ ml O_2_ /cm^2^ s) and (B) *J*_conv_/*J*_total_ as a function of *u*, at P_O2_= 30, 50, and 100 mmHg from right to left.

### 3.1 Convective and diffusive transport evaluated by *Pe*

i. **Convection regime** (*Pe* > 1) In the convection regime, *ub* >> *D*, exp(-*ub*/*D*) ~ 0, *J*_diff_ = *DM*_0_/*u*, and *J*_diff_ ~ 0. Thus, *P*_O2_ is constant across the tissue. Then, the O_2_ transport solely relies on *u* expressed as:

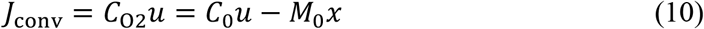 At *x* = 0, the O_2_ flux into the tissue is *J*_conv_ = *J*_total_ = *C*_0_*u*, with *J*_conv_/*J*_total_ = 1, while *J*_total_ depends on *C*_0_ and *u*.
ii. **Convection-diffusion regime** (*Pe* ~ 1) At *x* = 0, Eq. 5 becomes:

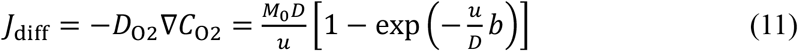

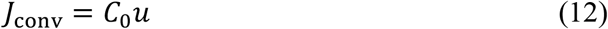

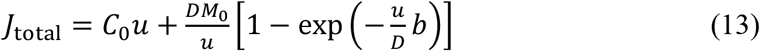 In this case, *J*_diff_/*J*_conv_ ~ *u*^−2^. ∇*C*_O2_ is sensitive to *u*. *J*_total_ increases with *u* while ∇*C*_O2_ decreases, resulting in a higher convective contribution. In the convection-diffusion regime, both *J*_diff_ and *J*_conv_ are dependent on *u*.
iii. **Diffusion**/**convection-diffusion regime** (*Pe* < 1) Normally, *Pe* < 1 implies a dominant O_2_ diffusion transport. However, the situation can be complicated by carefully considering the flow effects. In this regime, if *u* is large enough to modify Δ*P*_O2_, convection can also contribute significantly to *J*_total_ comparable to the contribution of diffusion.

#### Critical flow rate *u*_dc_

In this regime, *u* < *D*/*b*, 1 – exp(-*ub*/*D*) ≈ *ub*/*D*. Then, Eqs. 7-9 become:

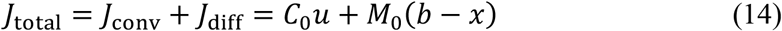

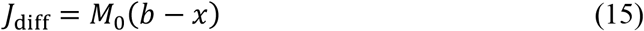

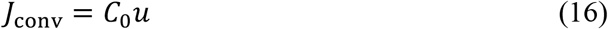

Set *x* = 0 in Eqs. 14-16, *J*_conv_/*J*_total_ can be expressed as a sigmoid function:

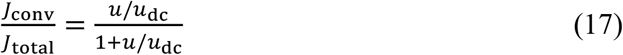

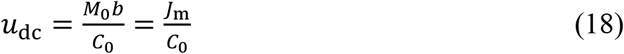

where, *u*_dc_ is introduced as a critical flow rate at which *J*_conv_ is equal to *J*_diff_. In its turn, *u*_dc_ is determined by the total tissue consumption *J*_m_ and *C*_0_.

#### Spatial boundary *x*_0_ for *J*_D_ = *J*_conv_

For *D*/*b* > *u* > *u*_dc_, *J*_total_ is governed by the convective transport, *J*_conv_ > *J*_diff_. The introduced *u*_dc_ gives an evaluation of the *J*_conv_ contribution to *J*_total_.

For *u* < *u*_dc_, *J*_total_ is dominated by the diffusive transport. At this condition, the contribution of *J*_conv_ and *J*_diff_ varies along *x*, due to the *J*_diff_ decrease with increasing *x* as shown in Fig. 2(D). The spatial boundary *x*_0_ is derived where *J*_diff_ = *J*_conv_ in Eqs. 15-16:

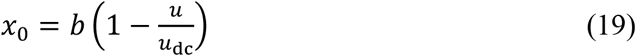

For *x* < *x*_0_, *J*_diff_ > *J*_conv_, the O_2_ transport to the site is prevalent by *J*_diff_. For *x* > *x*_0_, *J*_diff_ < *J*_conv_, the O_2_ transport to the site mainly relies on *J*_conv_. As a result, even for *u* < *u*_dc_, a *J*_conv_ dominated zone (*x* > *x*_0_) still exists.

Therefore, *x*_0_ can be used to evaluate the spatial contribution of the *J*_conv_ dominance for *u* < *u*_dc_. For example, if *u* = *u*_dc_, then *x*_0_ = 0 and the entire tissue satisfies *x* > *x*_0_, thereby *J*_diff_ < *J*_conv_ across the tissue. Hence, even for a small *u* across the tissue, the flow would dominate the O_2_ transport to the site far away from the O_2_ source.

### 3.2 O_2_ deficient tissue *J*_m_ < *bM*_0_

In the regime with *Pe* < 1, two feature parameters (*u*_dc_ and *x*_0_) are discussed above to evaluate the significance of the convection contribution of the fluid movement. As mentioned earlier, *L*_p_ is the length that only O_2_ diffusion can reach at *u* = 0. Hence, if *b* > *L*_p_, the O_2_ transport only by diffusion cannot supply O_2_ to the entire tissue and part of the tissue is O_2_ deficient without flow (*u* = 0). Flow would be key to delivery to the O_2_ deficient region. The expressions for *u*_dc_ and *x*_0_ should be modified by substituting *b* by *L*_p_ in Eqs. 18-19:

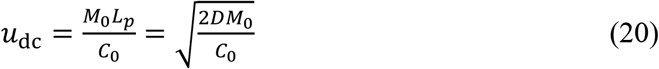

and

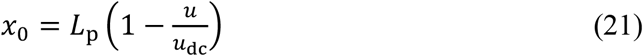

Fig. 4(A) shows that *u*_dc_ drops as *P*_0_ increases. *u*_dc_ can be calculated from Eq. 20 for hypoxia tissue *b* > *L*p and from Eq. 18 for *b* < *L*_p_. For the O_2_ deficient tissue, *u*_dc_ is also correlated to the O_2_ diffusion coefficient *D*.

**Fig. 4.**
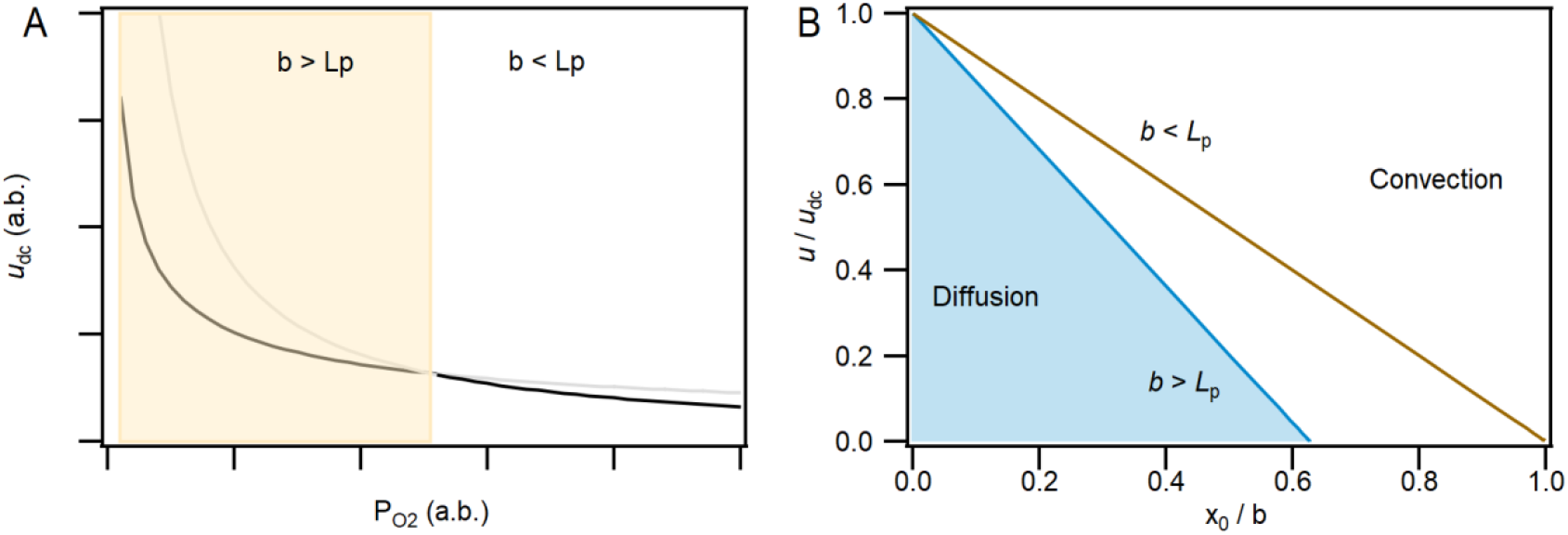
For *b* < *L*_p_ and *b* > *L*_p_, (A) *u*_dc_ as a function of *P*_O2_ (*J*_m_= 2× 10^−7^ ml O_2_ /cm^2^ s) and (B) the normalized *u* and *x*_0_;

Furthermore, *x*_0_ moves from the tissue scale (*x* = *b*) to the O_2_ source (*x* = 0) as *u* rises from 0 to *u*_dc_, as shown in Fig. 4(B). In the hypoxia condition, *i.e*., *b* > *L*_p_, ∇*C*_O_2__ cannot provide enough O_2_ reaching the tissue in the deep. Hence, the hypoxia region is sensitive to the fluid movement *u*.

### 3.3 Evaluation of *u_dc_* in tissue

The interspace is full of interstitial gel like matrix and fluid filtrated in and out the cell membrane. Even though the interstitial flow is neither uniform nor 100% laminar, *u*_dc_ can be used to evaluate the convection transport inside the tissue at certain sites. From Eqs. 18 and 20, *u*_dc_ is a function of *J*_m_ inside the tissue and *C*_0_ as shown in Fig. 5(A). *J*_m_ is the flux consumed by the tissue and its value is determined by *bM*_0_ for *b* < *L*_p_ and *L*_p_*M*_0_ for *b* > *L*_p_.

**Fig. 5.**
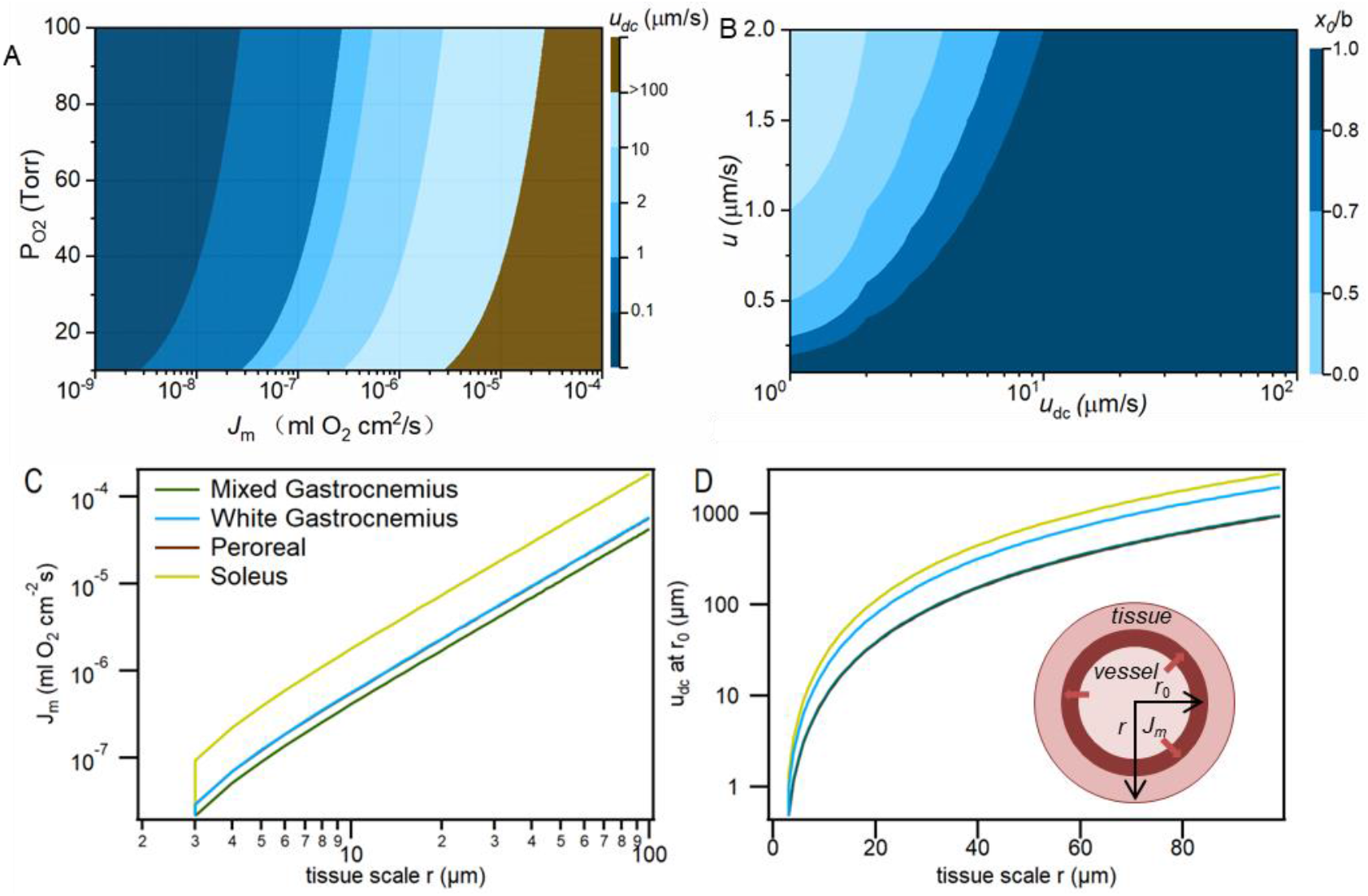
*u_dc_* as a function *J*_m_ and *P_0_*; (B) *x*_0_/b as the function of *u*_dc_ and the interstitial flow rate of *u* = 0.1 - 2 μm/s;(c) the tissue consumption flux *J*_m_ as the function of the tissue scale *r*; The *u*_dc_ as the function of the tissue diamete *r*.

Dissimilar *P*_O2_ in arteriolar, capillary, and venules have been reported for different tissues and organs such as brain, muscle, liver, *etc*. [39]. For instance, *P*_O2_ is in the range of 10 to 60 mmHg for capillary and 10 to 40 mmHg for tissue. The *P*_O2_ in arteriolar depends on the diameter and its range varies from 80 to 20 mmHg. A large drop across the vessel wall has been reported [39].

There are different types of fluid movement inside the tissue, including interstitial flow, perivascular fluid, and cerebrospinal fluid (CSF) in brain [40]. The CSF in cerebral aqueduct is reported as 5.27±1.77 cm/s [41]. From Fig. 5(A), CSF could delivery huge amounts of O_2_ to the brain tissue due to the large flow rates. Assuming *P*_0_ = 20 mmHg in cerebral aqueduct, *u*_dc_ = 5.27 cm/s, α = 3.3×10^−5^ ml·O_2_/(ml mmHg)[42], CSF can deliver a large consumption O_2_ flux to the tissue, *J*_m_ = *u*_dc_α*P*_0_ = 3.5×10^−3^ ml O_2_/cm^2^s. Hence, we predict that the large fluid movement of CSF can contribute significantly to the O_2_ delivery in the brain tissue. Another type of fluid movement inside the brain is the perivascular fluid around arterioles and venules, which plays a vital role in driving the clearance of amyloid-β (Aβ) in the brain. The typical average flow rate of the perivascular fluid is around 19 μm/s [43], which can also provide a large O_2_ flux to support the tissue consumption rate: *J*_m_ = 1.2× 10^−6^ ml O_2_/cm^2^ s (*P*_0_ = 20 mmHg). It suggests that the fluid movement would deliver large O_2_ flux away from the vessels.

#### Interstitial flow

The typical interstitial flow rate in tissue is in the range of 0.1 to 2 μm/s [18]. It can be seen from Fig. 5(A) that for the tissue consumption *J*_m_ < 2×10^−7^ ml O_2_/cm^2^s at *P*_O2_ = 40 mmHg, the interstitial flow dominates the O_2_ contribution to the tissue at *u* = 2 μm/s. The O_2_ flux measured from the microvessel to the tissue *in vivo* is around 10^−6^-10^−4^ ml O_2_/cm^2^s and also depends on the measurement and the diameter [11]. From Fig. 5(A), the corresponding *u*_dc_ range is 1 to 100 μm/s, which is much larger than the measured interstitial flow rate. However, Fig. 5(B) shows that there still exists a spatial region *x*_0_/b far away from the O_2_ source where pure diffusion based on ∇*P*_O2_ cannot meet *J*_m_. Although the input O_2_ flux for the whole tissue is dominated by ∇*P*_O2_, *i.e*., the diffusive transport, the interstitial flow can contribute significantly to the O_2_ flux to the far end region from the capillary source where ∇*P*_O2_ is small.

Table 1 lists *P*_O2_ in the interstitial space and the consumption rates for different types of the muscle tissue measured by the phosphorescence quenching method [9]. The experimental data in Table 1 can be used to evaluate the interstitial flow contribution in tissue. For the diameter of the vessel *r*_0_, the flux delivery through the interstitial flow outside the vessel to provide the tissue consumption is determined by *M*_0_*ρV*, where *V* is the tissue volume and *ρ* is the tissue density (1 g/cm^3^) as shown in the inset of Fig 5 (D). Then, *J*_m_ is a function of the tissue diameter *r*.

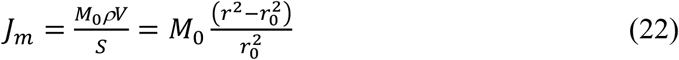

**Table 1.**
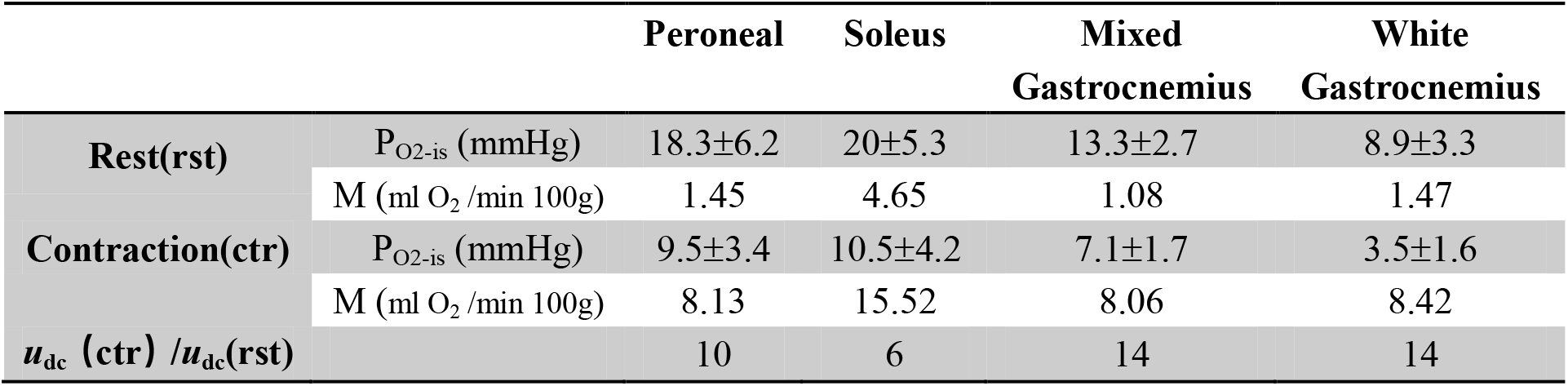
Consumption rate *M* and *Po_2_* in interstitial space for different muscles [9].

Fig. 5(C) shows *J*_m_ as a function of *r* for *r*_0_ = 2 μm, while Fig. 5(D) provides the corresponding *u*_dc_ at *r*_0_ near the vessel from Eq. 18. Here, *r* is the average distance for each capillary supply for the tissue consumption.

Inside the tissue, the heart has the highly density of the capillaries of 2500-4500 capillaries/mm^2^[44], which corresponds to the space of 12-22 μm between the capillaries. Hence, the corresponding *r* for the heart tissue is around 6-11 μm. For the skeletal muscles, the capillary density is 250 to 1500 /mm^2^. The reported capillary density for gastrocnemius and soleus is 365 and 288/mm [45], respectively, which corresponds to *r* of 25 and 29 μm. From Fig. 5(D), *u*_dc_ for the fluid movement around capillary is 60 μm/s for gastrocnemius and 2×10^2^ μm/s for soleus. Hence, in the hemostasis where the interstitial flow is 0.1-2 μm, the convective transport to the mitochondria near the capillary based on fluid movement can be negligible.

However, *u*_dc_ is dependent on *r*, *i.e*., the capillary density. For the experiments in Table 1, the capillary volume corresponds to 85% of the vascular volume [9]. If *r* is reduced to 5 μm, *u*_dc_ = 2 μm/s for gastrocnemius and 6 μm/s for soleus, the interstitial flow could contribute significantly to the O_2_ supply for the tissue. Moreover, if the tissue is under hypoxia with a reduced tissue consumption flux of *J*_m_ = 10^−7^ ml O_2_/cm^2^s, *u*_dc_ = 1.5 μm/s (20 mmHg) for soleus and 2.3 μm/s (13 mmHg) for mixed gastrocnemius. Therefore, the interstitial flow could contribute significantly to the site, which is far away from the capillary or the tissue under hypoxia conditions.

Table 1 also shows the consumption rates during the muscle contraction. The *P*O_2_ in the capillary drops significantly with a negligible change of Δ*P*_O2_ across the capillary wall but with a 3-6 fold enhancement of tissue consumption rates *M*_0_ [9]. *u*_dc_ under the contraction condition is around 6-14 times that at the rest. As reported, the interstitial flow can be enhanced by more than 10 times for the sports situation [1]. The microvascular permeability has been reported to associated with the openings in microvascular endothelium, which causes transcellular pathways for fluid [46]. This suggests that the interstitial flow could be enhanced during contraction, allowing for more fluid into the tissue, and bringing more O_2_ to the tissue.

#### Fluid movement inside myocyte

The intramyocyte Δ*P*_O_2__ is very low as reported by many groups [7]. Inside the cell, the fluid movement of cytoplasm depends on the cell size, which is significant in the plant, such as the cytoplasm streaming that can reach 0.1 mm/s. For the skeletal muscle cell, the fluid movement is much slower. However, inside muscle fibers, myoglobin (Mb) can store O_2_ and facilitate O_2_ transport at the same time [2]. Muscle type I fiber has the largest concentration of mitochondria and myoglobin, while muscle type IIX fiber has the lowest concentration. Hence, the cytoplasm inside the muscle could influence the myoglobin concentration redistribution and thereby the O_2_ delivery to the mitochondria inside the cell.

Mitochondria are mostly distributed around subsarcolemma and the deep-tissue mitochondria are reported to be interconnected and surrounded by myoglobin in the cytoplasm [47]. As a result, the myoglobin concentration is also included in the *u*_dc_ evaluation of the fluid movement inside myocyte. The concentration of the O_2_ loaded myoglobin (*C*_Mb-O2_) can be expressed as a function of *C*_O2_ according to:

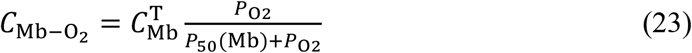

where, *C*_Mb_^T^ is the total concentration of uniformly distributed myoglobin (0-1 mM) [2] and *P*_50_(Mb) = 3 mmHg[6]. Substituting Eq. 22 into Eq. 1, the diffusive O_2_ flux inside the cell becomes:

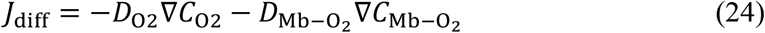

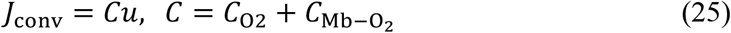

where, *D*_Mb_ (0.2×10^−6^ cm^2^/s) is the diffusion coefficient of myoglobin [48] by assuming *D*_Mb_ = *D*_Mb-O2_.

To evaluate the contribution of fluid movement inside the muscle, *C*_0_ in Eq. 18 becomes *C*_O2_ + *C*_Mb–O_2__. *u*_dc_ = *J*_m_/(*C*_O2_ + *C*_Mb–O_2__) can still be used as the criterion. *P*_O2_ of the intramyocyte is 20-30 mmHg [7]. *J*_m_ is determined by the mitochondrial density. For an isolated mitochondrion, the volume consumption *Γ* is about 1.3 ×10^−6^ mol/ml s [49]. Typically, mitochondrion is a prolate spheroid with the length of 4.3 μm and a diameter of 0.41 μm, yielding a volume *V*_m_ = 0.38 μm and surface area *S*_m_ = 4.4 μm for the dog hindlimb muscle. The consumption flux for an isolated mitochondrion can be estimated as *J*_m_ = *ΓV*_m_/*S*_m_ = 2×10^−7^ ml O_2_ /cm^2^ s. Fig. 6 shows *u*_dc_ as a function of *C*_mb_ for *P*_O2_ = 30 mmHg with *J*_m_ = 2×10^−7^ ml O_2_ /cm^2^ s.

**Fig. 6.**
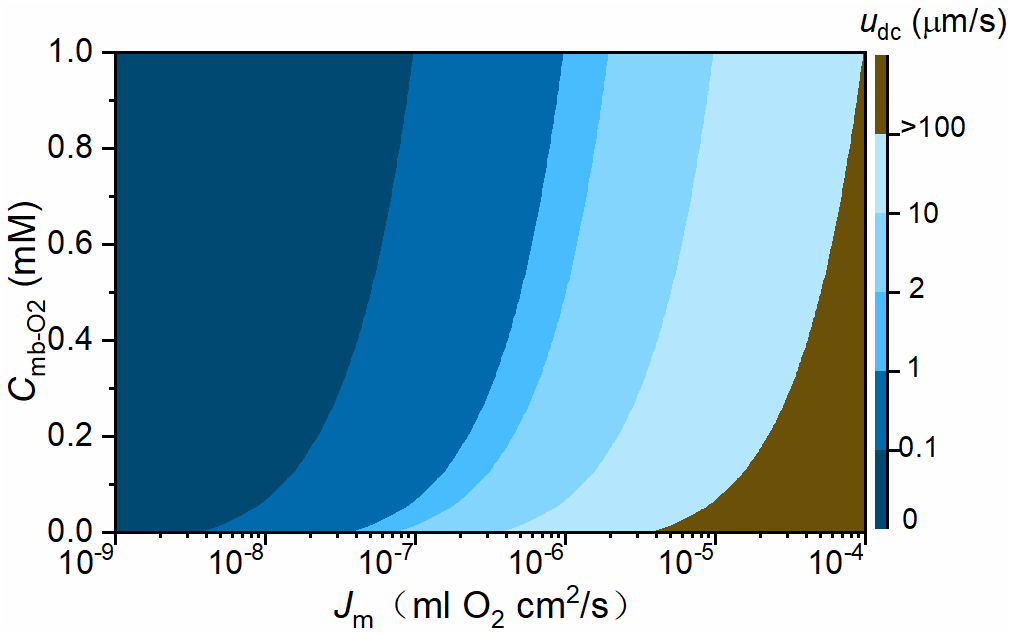
*u*_dc_ as a function of myoglobin concentration *C*_mb_ for P_O2_=30 mmHg with *J*_m_= 2 × 10^−7^ ml O_2_ /cm^2^ s for an isolated mitochondrion.

If *C*_mb_ increases from 0 to 0.5 mM, *u*_dc_ dramatically drops from 2 to 0.2 μm/s. It suggests that *C*_mb_ inside the tissue could also enhance the effects of the O_2_ transport contribution by the cytoplasm movement. For the muscle fiber with higher myoglobin concentrations, the higher cytoplasm movement rate would delivery more O_2_ to the mitochondrion.

### 3.4 Hydraulic pressure gradient and tissue permeability

The fluid movement inside the tissue is correlated to the tissue mobility and the local pressure gradient. The interstitial space is full of collagen fibers and proteins. Hence, the interstitial flow can be described by the porous media using Darcy’s law [50]. The flow is directly proportional to the hydraulic pressure gradient, ∇*P*_H2O_. *u* through the tissue can be expressed as follows:

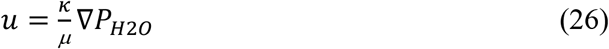

where, *κ* is the tissue permeability or specific hydraulic conductivity of tissue, in unit of cm^2^ and *μ* is the dynamic viscosity of the fluid. The value of *κ* depends on the tissue type, the hydrate of the interspace, pressure, and interstitium composites such as proteins, collagen fibers, *etc*. [50, 51]. For the interstitial flow, the composition varies by tissue locations. In most soft tissues, the interstitial flow has a composition similar to that of the blood plasma, with *μ* = 1.2 ×10^−3^ Pa s = 0.9×10^−5^ mmHg s [18].

The linear relationship between ∇*P*_H2O_ and *κ* is shown in Fig. 7, without considering the pressure dependence of *κ*. At the same pressure, the flow is much easier to go through the tissue with low *κ*. Typical *κ* values are in the range of 10^−7^ to 10^−11^ cm^2^ [18], while it can increase by more than 10^5^ times for *P*_H2O_ above the atmospheric pressure [51], *e.g*., in acute edema conditions. Therefore, the interstitial flow in tissue with acute edema is sensitive to the external force, which suggests that even a small force can induce a high flow rate and regulate the O_2_ transport inside the tissue. The external force such as massage, walk, sports, could also impact the pressure gradient in tissue.

**Fig. 7.**
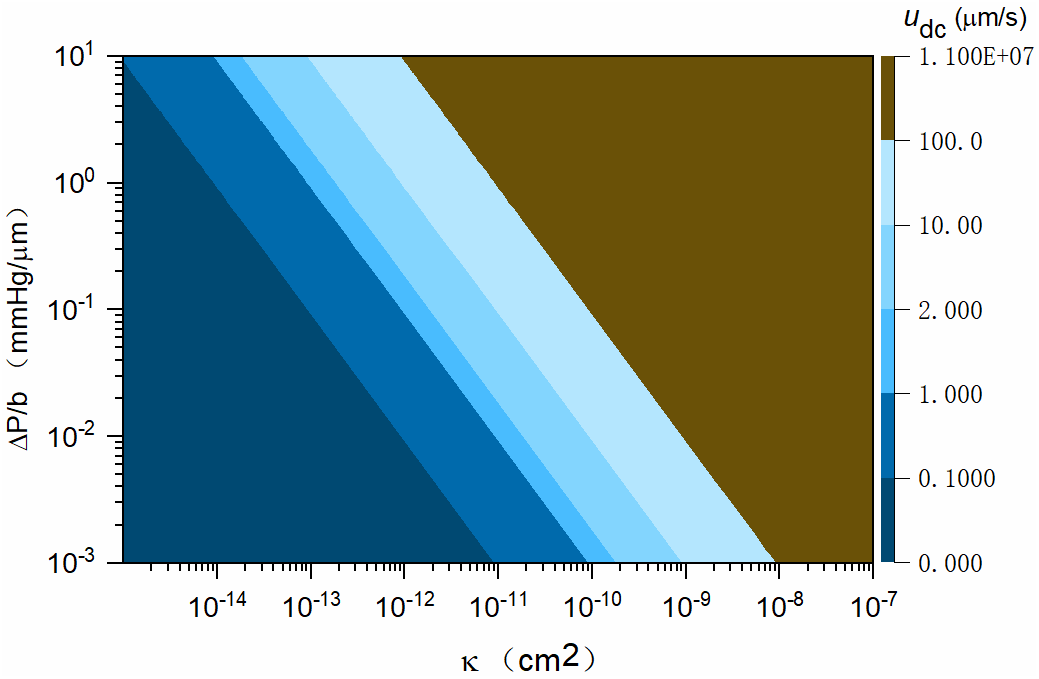
*u*_dc_ as the function of ∇*P_H2O_* and tissue permeability *κ*.

At the arterial end of a capillary, there is a net outward pressure of 10 mmHg including the osmotic pressure (due to protein concentration) and the hydrostatic pressure, which can push the fluid into the interstitial space and become re-adsorbed by venules. The average interstitial flow is directly *in vitro* measured as 0.6 μm/s with an estimated ∇*P*_H2O_ = 4 mmHg/μm in normal and neoplastic tissues [52].

Moreover, for myocyte, the muscle fibers with a high concentration of Mb, the O_2_ contribution of the cytosol movement is sensitive to ∇*P*_H2O_, which could accelerate the O_2_ transport inside the muscle tissue.

### 3.5 Discussion: Active and passive O_2_ transport inside tissue

Our analysis suggests that inside the body tissue, the O_2_ transport would have two mechanisms to support the tissue O_2_, diffusion and convection. Based on our analysis, we provide a complete picture of the of the active and passive O_2_ transport inside the tissue. The O_2_ flux inside the tissue has been modified to include two terms: diffusive transport (passive) and convective transport (active). Hydraulic pressure gradient is one of the important factors to drive the fluid movement and further contribute to the convective transport

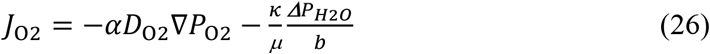

Hence, the O_2_ transport from the capillary to the tissue is affected by both ∇*P*_O2_ and ∇*P*_H2O_. The interstitial fluid movement may ensure the O_2_ flux supply for the far end of a mitochondrion from the capillary and is especially important for a damaged tissue where the diffusion supply cannot sustain the mitochondrion. The O_2_ transport can also be active by applying external pressure such as massage. Fig. 8 schematizes the O_2_ transport flux from capillary to mitochondria in tissue. Regulation of the blood vessel not only enhances *P*_O2_ in the capillary but also pump more plasma into the interstitial space to enhance the mass transport of O_2_ by vessel dilation.

**Fig. 8.**
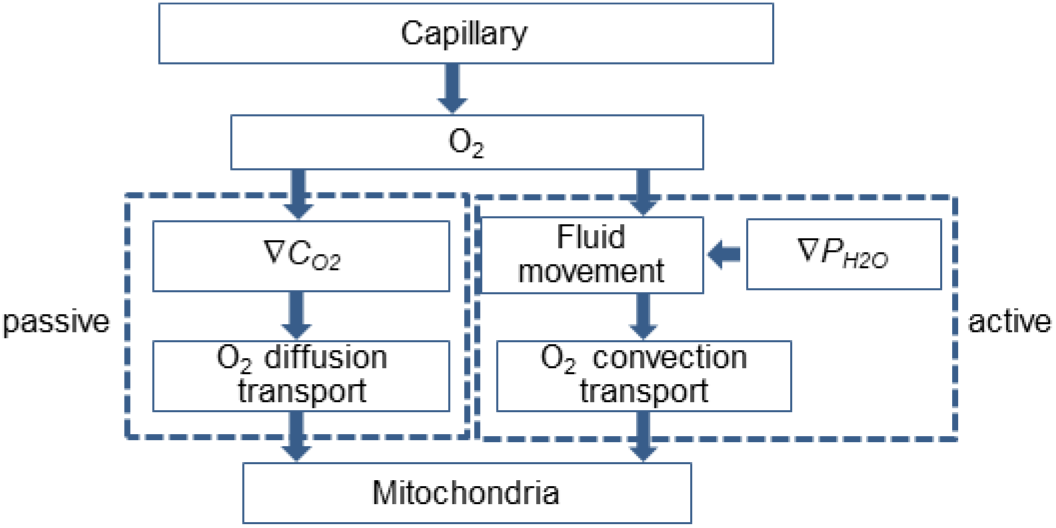
Summary of the active and passive O_2_ transport routes in tissue.

## 4. Conclusions

Krogh’s model is the foundation of the O_2_ transport through the passive diffusion in tissue. Normally, the fluid movement contribution to the O_2_ transport is analyzed based on *Pe*. As the analysis usually concerns small tissue scales with *Pe* < 1, the contribution of fluid movement to the O_2_ flux has long been ignored. In this work, we have shown that the O_2_ flux to sustain the mitochondrial consumption depends on not only the concentration gradient, but also the O_2_ flux contribution of fluid movement. The latter is significant for the region where the diminishing O_2_ gradient cannot sustain the needed O_2_ flux. Here, a comprehensive picture of O_2_ transport is presented by including the convective contribution from the fluid movement, in addition to the passive diffusion.

For tissue scales of *Pe* < 1, a criterion *u*_dc_ = *J*_m_/*C*_0_ is provided to evaluate the contribution of the fluid movement inside the tissue by a sigmoid function of *J*_conv_/*J*_total_ = (*u*/*u*_dc_)/(1+*u*/*u*_dc_). If the O_2_ flux in the local tissue and *u* of the fluid movement in the site are known, *u*_dc_ can be used to estimate the convective transport of O_2_ in tissue. For damaged tissues in particular, the convective contribution to the O_2_ transport can be more significant than diffusion.

The fluid movement including the cytoplasm and the interstitial flow offers an alternative active route for the O_2_ transport from capillary to tissue along with its counterpart of diffusion. It also provides a perspective to understand the morphogenesis and differentiation of cells. Moreover, this comprehensive picture of the O_2_ transport suggests that traditional therapies, such as massage and stretch, could facilitate the O_2_ transport from capillary to tissue so as to promote the tissue recovery from hypoxia. Hence, to understand the physiology of the O_2_ transport from the capillary to the tissue mitochondria, the convective contribution to the interstitial flow should be considered.

## Acknowledgement

This work was partially supported by the Chinese oversea program CSC foundation 201907035003 and the faculty fund of Uppsala University.

## Authors contributions

Z.L. conceived the idea. Z.L. discussed with C.W. on the physiology aspects of the idea and with S.-L.Z. on the physics aspects. Z.L. worked on model design, mathematical derivations, data analysis, and drafted the manuscript. C.W. built the COMSOL model. All authors contributed to the manuscript revision.

## Notes

### Competing Interest Statement

The authors have declared no competing interest.

### Summary of Updates

1. Updating the discussion and the title of the manuscript. 2. Fig. 1-4, 6-7 are revised, and Fig 5 is added, 3. added the udc evaluation for the interstitial flow and cytoplasm movement. 4. Updating the discussion and the conclusion

## References

1. Widmaier, E.P., H. Raff, and K.T. Strang, Vander’s Human Physiology: the mechanisms of body function. 2007: McGraw-Hill High Education.

2. Popel, A.S., Theory of oxygen transport to tissue. Critical reviews in biomedical engineering, 1989. 17(3): p. 257.

3. Krogh, A., The number and distribution of capillaries in muscles with calculations of the oxygen pressure head necessary for supplying the tissue. J Physiol, 1919. 52(6): p. 409–15.

4. Hill, A.V., The diffusion of oxygen and lactic acid through tissues. Proceedings of the Royal Society of London. Series B, Containing Papers of a Biological Character, 1928. 104(728): p. 39–96.

5. Poole, D.C., et al., August Krogh’s theory of muscle microvascular control and oxygen delivery: a paradigm shift based on new data. 2020. 598(20): p. 4473–4507.

6. Richardson, R.S., et al., Myoglobin O2 desaturation during exercise. Evidence of limited O2 transport. J Clin Invest, 1995. 96(4): p. 1916–26.

7. Poole, D.C., T.I. Musch, and T.D. Colburn, Oxygen flux from capillary to mitochondria: integration of contemporary discoveries. Eur J Appl Physiol, 2022. 122(1): p. 7–28.

8. Hirai, D.M., et al., Skeletal muscle microvascular and interstitial PO2 from rest to contractions. J Physiol, 2018. 596(5): p. 869–883.

9. Colburn, T.D., et al., Transcapillary PO(2) gradients in contracting muscles across the fibre type and oxidative continuum. J Physiol, 2020. 598(15): p. 3187–3202.

10. Popel, A.S., R.N. Pittman, and M.L. Ellsworth, Rate of oxygen loss from arterioles is an order of magnitude higher than expected. Am J Physiol, 1989. 256(3 Pt 2): p. H921–4.

11. Vadapalli, A., R.N. Pittman, and A.S. Popel, Estimating oxygen transport resistance of the microvascular wall. American Journal of Physiology-Heart and Circulatory Physiology, 2000. 279(2): p. H657–H671.

12. Pias, S.C., How does oxygen diffuse from capillaries to tissue mitochondria? Barriers and pathways. 2021. 599(6): p. 1769–1782.

13. Levich, V.G., Convective diffusion in liquid, in Physicochemical Hydrodynamics. 1962, Prentice-Hall Inc: Englewood Cliffs, N.J. p. 53.

14. Guyton, A.C. and J.E. Hall, The Microcirculation and the Lymphatic System: Capillary Fluid Exchange, Interstitial Fluid, and Lymph Flow, in Textbook of medical physiology. 2006, Elsvier Saunders. p. 184–193.

15. Skalak, T., G. Schmid-Schönbein, and B. Zweifach, New morphological evidence for a mechanism of lymph formation in skeletal muscle. Microvascular research, 1984. 28(1): p. 95–112.

16. Schmid-Schonbein, G.W., Microlymphatics and lymph flow. Physiological reviews, 1990. 70(4): p. 987–1028.

17. Mazzoni, M.C., T.C. Skalak, and G.W. Schmid-Schonbein, Effects of skeletal muscle fiber deformation on lymphatic volumes. American Journal of Physiology-Heart and Circulatory Physiology, 1990. 259(6): p. H1860–H1868.

18. Swartz, M.A. and M.E. Fleury, Interstitial flow and its effects in soft tissues. Annu. Rev. Biomed. Eng., 2007. 9: p. 229–256.

19. Havas, E., et al., Lymph flow dynamics in exercising human skeletal muscle as detected by scintography. The Journal of Physiology, 1997. 504(1): p. 233–239.

20. Swartz, M.A., The physiology of the lymphatic system. Advanced drug delivery reviews, 2001. 50(1-2): p. 3–20.

21. Aukland, K. and R.K. Reed, Interstitial-lymphatic mechanisms in the control of extracellular fluid volume. Physiological reviews, 1993. 73(1): p. 1–78.

22. Reddy, S.T., et al., A sensitive in vivo model for quantifying interstitial convective transport of injected macromolecules and nanoparticles. Journal of applied physiology, 2006. 101(4): p. 1162–1169.

23. Dukhin, S.S. and M.E. Labib, Convective diffusion of nanoparticles from the epithelial barrier toward regional lymph nodes. Advances in colloid and interface science, 2013. 199: p. 23–43.

24. Olszewski, W.L. and A. Engeset, Intrinsic contractility of prenodal lymph vessels and lymph flow in human leg. American Journal of Physiology-Heart and Circulatory Physiology, 1980. 239(6): p. H775–H783.

25. McGuire, S., D. Zaharoff, and F. Yuan, Nonlinear dependence of hydraulic conductivity on tissue deformation during intratumoral infusion. Annals of biomedical engineering, 2006. 34(7): p. 1173–1181.

26. Zhang, X.-Y., et al., Interstitial hydraulic conductivity in a fibrosarcoma. American Journal of Physiology-Heart and Circulatory Physiology, 2000. 279(6): p. H2726–H2734.

27. Dreher, M.R., et al., Tumor vascular permeability, accumulation, and penetration of macromolecular drug carriers. Journal of the National Cancer Institute, 2006. 98(5): p. 335–344.

28. Heldin, C.-H., et al., High interstitial fluid pressure—an obstacle in cancer therapy. Nature Reviews Cancer, 2004. 4(10): p. 806–813.

29. Evans, R.C. and T.M. Quinn, Solute convection in dynamically compressed cartilage. Journal of biomechanics, 2006. 39(6): p. 1048–1055.

30. Hadida, M. and D. Marchat, Strategy for achieving standardized bone models. Biotechnology and Bioengineering, 2020. 117(1): p. 251–271.

31. Goldman, J., et al., Regulation of lymphatic capillary regeneration by interstitial flow in skin. American Journal of Physiology-Heart and Circulatory Physiology, 2007. 292(5): p. H2176–H2183.

32. Fleury, M.E., K.C. Boardman, and M.A. Swartz, Autologous morphogen gradients by subtle interstitial flow and matrix interactions. Biophysical journal, 2006. 91(1): p. 113–121.

33. Rutkowski, J.M. and M.A. Swartz, A driving force for change: interstitial flow as a morphoregulator. Trends in Cell Biology, 2007. 17(1): p. 44–50.

34. Li, H., et al., Active interfacial dynamic transport of fluid in a network of fibrous connective tissues throughout the whole body. Cell Proliferation, 2020. 53(2): p. e12760.

35. Zhang, W.-B., G.-J. Wang, and K. Fuxe, Classic and modern meridian studies: A review of low hydraulic resistance channels along meridians and their relevance for therapeutic effects in Traditional Chinese Medicine. Evidence-based Complementary and Alternative Medicine, 2015. 2015.

36. Ray, L.A. and J.J. Heys, Fluid Flow and Mass Transport in Brain Tissue. Fluids, 2019. 4(4): p. 196.

37. Cenaj, O., et al., Evidence for continuity of interstitial spaces across tissue and organ boundaries in humans. Communications Biology, 2021. 4(1): p. 436.

38. Dixon, J.B., et al., Lymph flow, shear stress, and lymphocyte velocity in rat mesenteric prenodal lymphatics. Microcirculation, 2006. 13(7): p. 597–610.

39. Pittman, R.N., Oxygen gradients in the microcirculation. Acta Physiologica, 2011. 202(3): p. 311–322.

40. Stine, C.A. and J.M. Munson, Convection-Enhanced Delivery: Connection to and Impact of Interstitial Fluid Flow. Frontiers in Oncology, 2019. 9.

41. Mase, M., et al. Quantitative Analysis of CSF Flow Dynamics using MRI in Normal Pressure Hydrocephalus. in Intracranial Pressure and Neuromonitoring in Brain Injury. 1998. Vienna: Springer Vienna.

42. Dutta, A., et al., Oxygen Tension Profiles in Isolated Hamster Retractor Muscle at Different Temperatures. Microvascular Research, 1996. 51(3): p. 288–302.

43. Mestre, H., et al., Flow of cerebrospinal fluid is driven by arterial pulsations and is reduced in hypertension. Nature Communications, 2018. 9(1): p. 4878.

44. Glancy, B. and R.S. Balaban, Energy metabolism design of the striated muscle cell. Physiological Reviews, 2021. 101(4): p. 1561–1607.

45. Andersen, P. and A.J. Kroese, Capillary supply in soleus and gastrocnemius muscles of man. Pflügers Archiv, 1978. 375(3): p. 245–249.

46. Michel, C.C. and C.R. Neal, Openings Through Endothelial Cells Associated with Increased Microvascular Permeability. Microcirculation, 1999. 6(1): p. 45–54.

47. Glancy, B., et al., Mitochondrial reticulum for cellular energy distribution in muscle. Nature, 2015. 523(7562): p. 617–620.

48. Covell, D.G. and J.A. Jacquez, Does myoglobin contribute significantly to diffusion of oxygen in red skeletal muscle? American Journal of Physiology-Regulatory, Integrative and Comparative Physiology, 1987. 252(2): p. R341–R347.

49. Clark, A., Jr. and P.A. Clark, Local oxygen gradients near isolated mitochondria. Biophys J, 1985. 48(6): p. 931–8.

50. Levick, J., Flow through interstitium and other fibrous matrices. Quarterly Journal of Experimental Physiology: Translation and Integration, 1987. 72(4): p. 409–437.

51. Guyton, A.C., K. Scheel, and D. Murphree, Interstitial fluid pressure: III. Its effect on resistance to tissue fluid mobility. Circulation Research, 1966. 19(2): p. 412–419.

52. Chary, S.R. and R.K. Jain, Direct measurement of interstitial convection and diffusion of albumin in normal and neoplastic tissues by fluorescence photobleaching. Proc Natl Acad Sci U S A, 1989. 86(14): p. 5385–9.

