## supplemental for "Evaluation of the fluid-movement contribution to the oxygen transport in tissue"

**Supporting information**

**Appendix**

**1. $O_2$ diffusive transport**

The tissue b has a constant $O_2$ consumption rate $M_0$ by the mitochondria. From Fick's first law, without considering convection contribution, the $O_2$ flux per area per second is:

$$J_{\mathrm{diff}} = -D_{\mathrm{O2}} \nabla C_{\mathrm{O2}} \quad \text{(A1)}$$

The flux into the tissue per second can be expressed by Fick's second law:

$$\frac{dC_{\mathrm{O2}}}{dt} = D_{\mathrm{O2}} \nabla^2 C_{\mathrm{O2}} - M_0 \quad \text{(A2)}$$

Eqs. A1 − A2 can be solved at the steady state $dC_{\mathrm{O2}}/dt = 0$ with the boundary condition: $C_{\mathrm{O2}} = C_0 = \alpha P_0$, at $x = 0$, and $\nabla C_{\mathrm{O2}} = 0$, at $x = b$.

$$C_{\mathrm{O2}} = C_0 + \frac{M_0}{2D} x(x - 2b) \quad \text{(A3)}$$

From Eq. A3, the penetration length $L_{\mathrm{p}} = (2DC_0/M_0)^{1/2}$ is obtained at $C_{\mathrm{O2}} = 0$. The total $O_2$ flux $J_{\mathrm{total}}$ into the tissue from the capillary corresponds to the $O_2$ flux at $x = 0$. Substituting Eq. A3 into Eq. A1, we can obtain:

$$J_{\mathrm{diff}} = M_0(b - x) \quad \text{(A4)}$$

At $x = 0$, $J_{\mathrm{diff}} = M_0 b$.

**2. Including $O_2$ convective transport**

If the total flux is determined by the gradient of concentration and the body fluid flow $\boldsymbol{u}$, the total $O_2$ flux becomes,

$$J_{\mathrm{total}} = J_{\mathrm{diff}} + J_{\mathrm{conv}} = -D_{\mathrm{O2}} \nabla C_{\mathrm{O2}} + C_{\mathrm{O2}} u \quad \text{(A5)}$$

$$\frac{dC_{\mathrm{O2}}}{dt} = D_{\mathrm{O2}} \nabla^2 C_{\mathrm{O2}} - \boldsymbol{u} \nabla C_{\mathrm{O2}} - M_0 \quad \text{(A6)}$$

The liquid is impressible and $\nabla \cdot \boldsymbol{u} = 0$. At the steady state and with the boundary conditions $C_{\mathrm{O2}} = C_0 = \alpha P_0$ at $x = 0$, and $\nabla C_{\mathrm{O2}} = 0$ at $x = b$, the equation can be solved. Let,

$$y = \frac{dC}{dx} + \frac{M_0}{u} \quad \text{(A7)}$$

Eq. A6 becomes:

$$0 = \mathrm{D} \nabla y - uy \quad \text{(A8)}$$

$$y = A \exp\left(\frac{u}{D} x\right) \quad \text{(A9)}$$

where, $A$ is a constant. With the boundary condition $\nabla C = 0$ at $x = b$, we can obtain

$$A = \frac{M_0}{u} \exp\left(-\frac{u}{D} b\right) \quad \text{(A10)}$$

$$y = \frac{M_0}{u} \exp\left[-\frac{u}{D}(b - x)\right] \tag{A11}$$

Substituting Eq. A11 into Eq. A7,

$$\frac{M_0}{u} \exp\left[-\frac{u}{D}(b - x)\right] = \frac{dC}{dx} + \frac{M_0}{u} \tag{A12}$$

$$C = \frac{M_0}{u}\left\{-x + \frac{D}{u}\exp\left[-\frac{u}{D}(b - x)\right]\right\} + B \tag{A13}$$

The constant *B* can be determined by the boundary condition $C_{O_2} = C_0$ at $x = 0$.

$$B = C_0 - \frac{DM_0}{u^2}\exp\left(-\frac{u}{D}b\right) \tag{A14}$$

Thus, Eq. A13 becomes:

$$C = C_0 - \frac{M_0}{u}x + \frac{DM_0}{u^2}\exp\left(-\frac{u}{D}b\right)\left[\exp\left(\frac{u}{D}x\right) - 1\right] \tag{A15}$$

The diffusion flux component is:

$$J_{\text{diff}} = -D_{O2}\nabla C_{O2} = \frac{M_0 D}{u}\left\{1 - \exp\left[-\frac{u}{D}(b - x)\right]\right\} \tag{A16}$$

The convection flux component is:

$$J_{\text{conv}} = C_0 u - M_0 x + \frac{DM_0}{u}\exp\left(-\frac{u}{D}b\right)\left[\exp\left(\frac{u}{D}x\right) - 1\right] \tag{A17}$$

The total flux is:

$$J = J_{\text{conv}} + J_{\text{diff}} = C_0 u + \frac{DM_0}{u}\left[1 - \exp\left(-\frac{u}{D}b\right)\right] - M_0 x \tag{A18}$$

If $ub < D$, following the first order expansion,

$$1 - \exp\left(-\frac{ub}{D}\right) = 1 - \left(1 - \frac{ub}{D}\right) = \frac{ub}{D} \tag{A19}$$

$$C = C_0 - \frac{M_0}{u}x - \frac{DM_0}{u^2}\exp\left(-\frac{u}{D}b\right)\left[1 - \exp\left(\frac{u}{D}x\right)\right]$$

$$C = C_0 - \frac{M_0}{u}x - \frac{DM_0}{u^2}\left(1 - \frac{ub}{D}\right)\left[1 - \exp\left(\frac{u}{D}x\right)\right]$$

$$C = C_0 - \frac{M_0}{u}x - \frac{DM_0}{u^2}\left[1 - \exp\left(\frac{u}{D}x\right)\right] + \frac{bM_0}{u}\left[1 - \exp\left(\frac{u}{D}x\right)\right] \tag{A20}$$

$$\left\{\exp\left(-\frac{u}{D}b\right) - \exp\left[-\frac{u}{D}(b - x)\right]\left(-\frac{u}{D}(b - x)\right)\right\} = \left(1 - \frac{ub}{D}\right) - \left[1 - \frac{u}{D}(b - x)\right] = -\frac{ux}{D}$$

(A21)

$$J = J_{\text{conv}} + J_{\text{diff}} = C_0 u + \frac{DM_0}{u}\left(\frac{u}{D}b\right) - M_0 x = C_0 u + M_0(b - x) \tag{A22}$$

$$J_{\text{diff}} = M_0(b - x) \tag{A23}$$

At $x = 0$,

$$J_{\text{total}} = C_0 u + \frac{DM_0}{u}\left[1 - \exp\left(-\frac{u}{D}b\right)\right] \tag{A24}$$

$$J_{\text{conv}} = C_0 u \tag{A25}$$

$$J_{\text{diff}} = \frac{M_0 D}{u}\left[1 - \exp\left(-\frac{u}{D}b\right)\right] \tag{A26}$$

For $ub < D$, $J_{\text{diff}} = M_0 b$.

$$\frac{J_{\text{diff}}}{J_{\text{conv}}} = \frac{M_0 b}{C_0 u} \tag{A27}$$
